# Chromosome-level genome assembly of the Erythrina Gall Wasp, *Quadrastichus erythrinae* (Hymenoptera: Eulophidae)

**DOI:** 10.64898/2026.01.23.701414

**Authors:** Y. Miles Zhang, Justin Merondun, Renee L. Corpuz, Angela N. Kauwe, Scott M. Geib, Sheina B. Sim

## Abstract

The erythrina gall wasp, *Quadrastichus erythrinae*, is an invasive gall-inducing chalcidoid wasp and a major pest of the endemic wiliwili tree (*Erythrina sandwicensis*) in Hawai□i. As a foundation to associated research, we generated a chromosome-level genome assembly from a wild-collected female measuring <2 mm. The final assembly consists of five scaffolds representing the five autosomes totaling 399 Mb (N50 = 75.6 Mb) and one unplaced 16 kb contig. BUSCO analysis recovers 91.1% of conserved Hymenoptera orthologs, representing the first chromosome-scale genome for the genus *Quadrastichus*. Comparative genomic analyses reveal syntenic conservation across Hymenoptera despite deep evolutionary divergence, with strongest collinearity to the chalcidoid *Nasonia vitripennis*. Genome size variation is largely explained by repeat content, and *Q. erythrinae* exhibits high proportions of unclassified transposable elements similar to the cynipid gall inducer. We also assembled a complete genome of its endosymbiont, *Wolbachia pipientis*. Together, these genomic resources provide a foundation for comparative, evolutionary, and applied research aimed at managing this invasive pest.

## Introduction

The tetrastichine eulophids (Chalcidoidea, Eulophidae, Tetrastichinae) are one of the largest subfamilies of chalcidoid wasps, comprising over 100 genera and approximately 3,000 known species (UCD Community, 2023). This hyperdiverse but poorly defined group mostly consists of parasitoids, but also includes 16 genera with recorded phytophagy and the ability to induce galls on both monocot and dicot plants (Zhang et al., 2022). *Quadrastichus* is one of the larger genera within the “*Tetrastichus* group” *sensu* Rasplus et al. (2020), with 95 described species and many others still undescribed (UCD Community, 2023; Kärnnäs et al., 2025).

The Erythrina gall wasp, *Quadrastichus erythrinae*, is a gall-inducing species first identified in 2004 when it emerged as a serious invasive pest (Kim et al., 2004). Despite being less than 2 mm in size, the wasp induces galls on the young leaves, stems, and petioles of *Erythrina* (Fabaceae: Faboideae) species (**Figure 1**), often leading to tree decline and death (Kim et al., 2004; Lin et al., 2021). Likely originating in East Africa, *Q. erythrinae* has spread rapidly across tropical and subtropical regions of Asia, Oceania, and the Americas (Kim et al., 2004; Lin et al., 2021). It causes significant damage to both endemic and introduced *Erythrina* species. *Qudrastistichus erythrinae* posed a major threat to *Erythrina sandwicensis* (wiliwili), the only endemic *Erythrina* species on the Hawaiian Islands. This tree holds great societal and cultural value for native Hawaiians and is of significant ecological importance as one of the few deciduous trees in the endangered lowland dry forests (Kaufman et al., 2020). The wiliwili was near the brink of extinction due to *Q. erythrinae* infestations, until the successful deployment of the biological control agent *Eurytoma erythrinae* reduced infestation levels (Kaufman et al., 2020).

**Figure 1.**
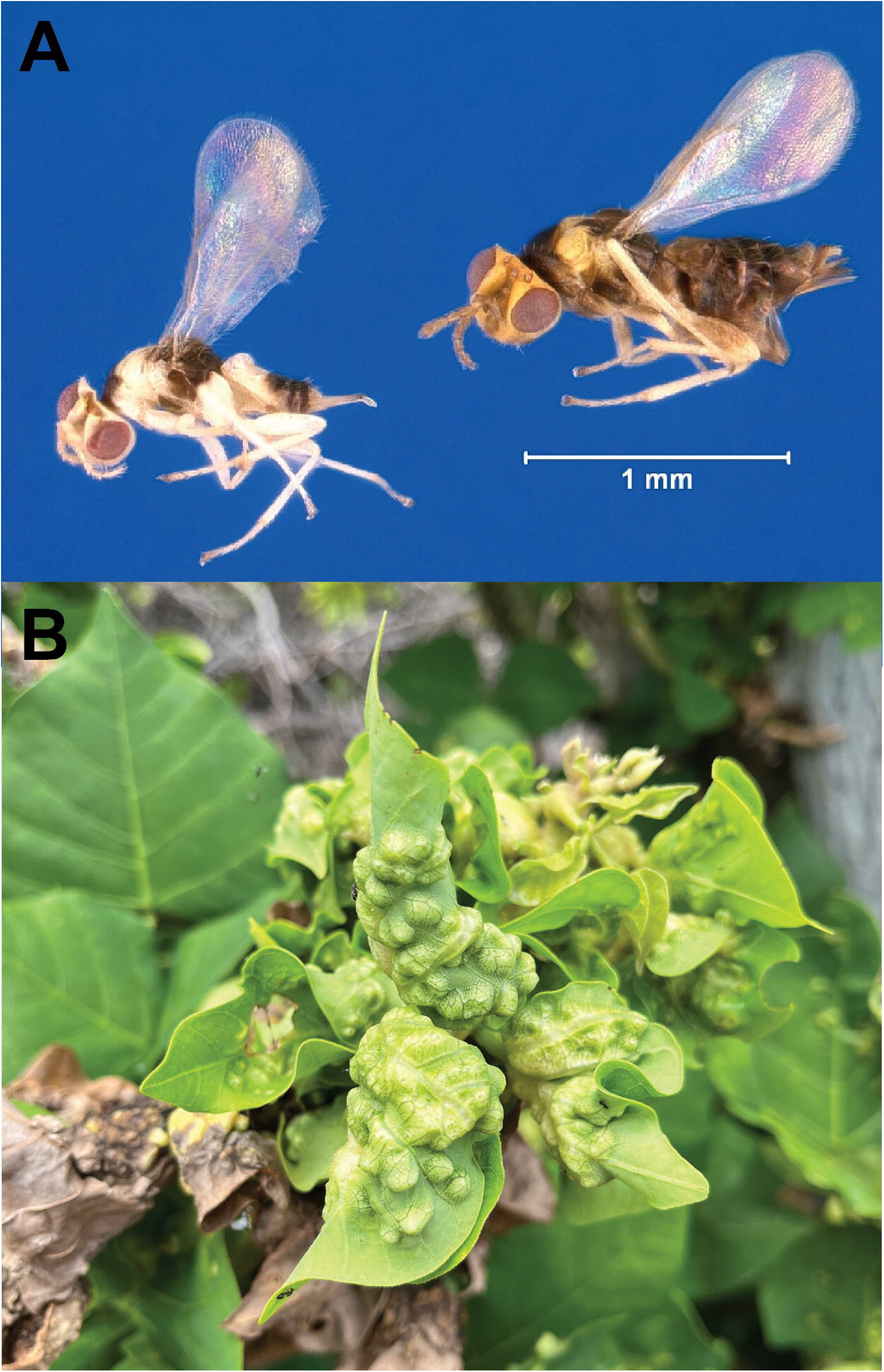
*Quadrastichus erythrinae*. A) Lateral habitus of male (left) and female (right) adult *Q. erythrinae*. Photo by Erich G. Vallery, USDA Forest Service – SRS-4552, Bugwood.org, licensed under CC BY 3.0. B) Galls induced by *Q. erythrinae* on wiliwili tree in Kona, Hawai□i.

We sequenced and assembled a chromosome-level genome for *Q. erythrinae*, leveraging its raw sequencing data to also recover the complete genomes of its *Wolbachia* endosymbionts. This approach demonstrates how mining host genome data can effectively characterize cobionts, providing crucial ecological context for their hosts.

## Materials and methods

### Source material

Galls from the leaves of *E. sandwicensis* were collected near Wawaloli Beach Park in Kaiminani, HI (19.7145, -156.0491) on 3.V.2025 and returned to the United States Daniel K Inouye U.S. Pacific Basin Agricultural Research Center in Hilo, Hawai□i, USA. Adults were reared out of the galls, identified, flash-frozen alive using liquid nitrogen, and stored at -80°C until genomic library preparation.

### Library preparation and sequencing

The whole body of a single female wasp was homogenized into a fine powder while kept frozen using a SPEX SamplePrep 2010 Geno/Grinder (Cole Parmer, Metuchen, New Jersey, USA) and underwent high molecular weight (HMW) DNA extraction using the fresh or frozen tissue protocol of the Qiagen MagAttract HMW DNA Kit (Qiagen, Hilden, Germany). Following isolation, the HMW DNA (92.5ng) was sheared to a mean fragment length of 20 Kb with a Megaruptor 3 (Diagenode, Dennville, New Jersey, USA) and prepared into a PacBio SMRTBell library using the SMRTBell Express Template Prep Kit. 3.0 (Pacific Biosciences, Menlo Park, California, USA) using a barcoded adapter. After DNA isolation, shearing, and library preparation, the sample was purified using solid-phase reversible immobilization beads (SPRI beads) (DeAngelis et al., 1995) and quantified using fluorometry and spectrophotometric absorbance ratios (DeNovix Inc., Wilmington, Delaware, USA). Fragment length distribution after each step were determined by Femto Pulse or Fragment Analyzer (Agilent Technologies, Santa Clara, California, USA). The resulting library was pooled with other samples and sequenced on a PacBio Revio system using a 30-hour movie length on 1/10th of a Revio SMRTCell. Raw subreads were converted to HiFi data using the PacBio SMRTLink software v.10.1.

Concurrent to HiFi sequencing, a pool of twelve males were used to prepare an enriched chromosome conformation capture (HiC) library. Tissues were homogenized in 1× phosphate buffered saline and nuclei were crosslinked in a 2% formaldehyde solution. Following crosslinking, the sample was lysed and digested using the restriction enzymes DdeI and DpnII. To enrich the sample for proximity ligated fragments, a biotin-labeled dATP fill-in step was performed prior to proximity ligation so fragments could be captured downstream. Following proximity ligation, a crosslink reversal step was performed followed by two DNA purification steps using SPRI beads, the removal of biotin from unligated ends, and another DNA purification step. The sample was size-selected using SPRI beads, biotinylated ligation products were captured, and the sample was prepared into a short-read sequencing library using the NEBNext Ultra II DNA Library Prep Kit (New England Biolabs, Ipswich, Massachusetts, USA). The final libraries were sequenced on a partial flowcell using the AVITI 2×150 Sequencing Kit Cloudbreak FS High Output kit on the Element AVITI System (Element Biosciences, San Diego, CA). Following sequencing, raw reads were base called using bases2fastq v.2.3.0.2116803307.

### Genome assembly, assessment, and contaminant removal

HiFi reads were screened and filtered for adapter-contaminated sequence artifacts using FCS-Adaptor and HiFiAdapterFilt (Sim et al., 2022) (https://github.com/ncbi/fcs). The resulting HiFi reads were used to assemble contigs using HiFiASM (v.0.24.0-r702) (Cheng et al., 2021; Cheng et al., 2022). The contig assemblies were subsequently purged of duplicate contigs using PurgeDups (Guan et al., 2020), and the duplicate purged contig assembly served as the reference to map HiC reads using BWAmem 2 (v.2.2.1) (Vasimuddin et al, 2019). The resulting mapped reads were filtered for artifact PCR duplicates using Picard (v.3.2.0) (Picard2019toolkit, 2019 https://github.com/broadinstitute/picard). A contact map was generated from the de-duplicated mapped reads using the YaHS pipeline (Zhou et al., 2023). Visualization of the contact map and minor manual editing was achieved using Juicebox (v.2.15) (Durand et al., 2016). Minimap2 (v.2.22-r1101) was used to map HiFi reads back to the contig assembly and calculate coverage of each contig, the –auto function and –genome mode of BUSCO (v.5.8.3, Manni et al. 2021) was used to select the appropriate taxon database and estimate genome completeness, and BLAST+ and Diamond were used to perform nucleotide alignments to the NCBI nucleotide database (accessed November 2025) and UniProt protein database (accessed November 2025) respectively (Buchfink et al., 2021; Camacho et al., 2009; Li, 2018, Tegenfeldt et al., 2025). The resulting outputs of minimap2, BUSCO, BLAST+, and Diamond were summarized and visualized using Blobtools2 and blobblurb (Challis et al., 2020); (https://github.com/sheinasim/blobblurb). Additional taxonomic assignments of contigs and contig fragments was performed using the FCS-GX (Astashyn et al., 2024). The confluence of the taxonomic assignments based on nucleotide alignment, protein alignment, and FCS-GX was used to identify contigs assigned to *Wolbachia* and remove non-Arthropod contigs from the assembly. The primary and alternate assemblies were submitted to the National Center for Biotechnology Information (NCBI).

### RNA extraction and sequencing

We sequenced RNA to aid in genome annotation. Specifically, total RNA was extracted from three pools of four male specimens separated by body parts (head, thorax/mesosoma, and abdomen/metasoma), through homogenization of snap frozen tissue in Tri Reagents and then extraction using the Zymo Direct-zol Magbead Total RNA kit (Zymo Research, Irvine, CA) on a Kingfisher Flex 96 system (ThermoFisher, Waltham, MA). From total RNA, a poly(A) cDNA library was generated using the NEB Ultra II RNA Library Prep Kit (New England Biolabs, Ipswich, MA) using oligo(dT) selection. The final RNASeq libraries were sequenced on a partial flowcell using the AVITI 2×150 Sequencing Kit Cloudbreak FS High Output kit on the Element AVITI System (Element Biosciences, San Diego, CA). Following sequencing, raw reads were base called using bases2fastq v.2.3.0.2116803307 the same way as HiC above.

### Genome Annotation

We performed genome annotation of *Q. erythrinae* using the Eukaryotic Genome Annotation Pipeline - External (EGAPx) (https://github.com/ncbi/egapx) v0.4.0. Short-read RNA-Seq from a pool of 4 whole male bodies were used as transcriptome evidence and aligned to the respective reference using STAR. Miniprot was used to align Hymenoptera protein sequences to the reference. Gnomon (https://www.ncbi.nlm.nih.gov/refseq/annotation_euk/gnomon/) was used for gene prediction using protein and RNA-seq alignments and *ab-initio* predictions based on HMM. Lastly EGAPx adds functional annotations to the final structural annotation set based on orthology and model type and quality. The *Wolbachia* genome was annotated using Prokka v1.14.6 (Seeman, 2014).

### Mitochondrial genome

We identified all potential mitochondrial genome contigs using the MitoHiFi pipeline (Uliano-Silva et al., 2023). MitoHiFi implemented a BLAST search for contigs that have a high similarity to the mitochondrial genome of *Nasonia vitripennis*, (NCBI RefSeq accession NC_066201.1) (Camacho et al., 2009) and selected the contig with the greatest similarity. Mitochondrial genes were then structurally annotated using intervals from the same mitochondrial genome used in the BLAST search through the MitFi annotation program in MITOS2 (Bernt et al., 2013). A representative mitochondrial genome was submitted to NCBI under the accession PX622211.

### Synteny across Hymenoptera

Synteny conservation among eight chromosome-scale diverse hymenopteran assemblies (Table 2) was used to place the *Q. erythrinae* genome in a comparative genomic context. Analyses were performed using all annotated genes by first extracting the longest isoform per gene from the RefSeq GFF annotations using AGAT (v1.4.3) (Dainat et al. 2025), with corresponding nucleotide CDS sequences inferred with TransDecoder (v5.7.1) (Haas et al. 2013). Synteny inference was performed in JCVI (v1.4.16) (Tang et al. 2024) using LAST (v2.37.4) (Frith et al. 2010) for pairwise alignment, applying a cscore threshold of 0.99 to retain reciprocal best-hit gene pairs and filtering syntenic blocks containing at least 10 gene anchors. Synteny was visualized using pairwise dot plots and whole-chromosome karyoplots. Note that JCVI visualizations are based solely on gene order and anchor counts, so chromosome lengths in JCVI plots do not reflect physical chromosome sizes.

We corroborated this analysis with BUSCO-defined genomic anchors and chromsyn (v1.6.1) (Edwards et al. 2022), which infers collinear regions from the relative order and spacing of conserved single-copy orthologs. All genomes were annotated with BUSCO (v5.8.3) using the Hymenoptera lineage dataset (hymenoptera_odb12). Conserved regions were defined by clustering BUSCO loci using chromsyn, retaining only syntenic blocks larger than 100 Kb which contain more than 2 BUSCO genes to emphasize macro-syntenic structure. Shared synteny among assemblies was quantified by counting these pairwise BUSCO-defined blocks with visualization using log-scaled heatmaps in R (v4.4.1) and the tidyverse (v2.0.0) (R Core Team 2017; Wickham et al. 2019).

### Repeat landscape

We quantified total repetitive DNA content for the same eight chromosome-level hymenopteran assemblies using EarlGrey (v6.3.5) (Baril et al. 2024). Repeat annotation summaries were derived from GFF files using custom R scripts, assigning each genomic position to a single repeat subclass using a fixed hierarchical order (LTR, LINE, Penelope, SINE, DNA, Rolling Circle, Other, Unclassified, Non-Repeat), with overlapping intervals resolved by priority-based merging in this order using R, tidyverse, and GenomicRanges (v1.56.2) (Lawrence et al. 2013). From these repeat annotations we extracted three summary metrics per transposable element (TE) subclass: (i) total cumulative genomic coverage (Mb), (ii) the number of distinct TE families, and (iii) the proportion of the genome covered by each subclass. To assess the relationship between repeat abundance and genome size while accounting for shared evolutionary history, we performed phylogenetic generalized least squares (PGLS) regression using caper (v1.0.4) (Orme et al. 2025) in R. Both genome size and total repeat genomic coverage (Mb) were log-transformed and Pagel’s λ was estimated by maximum likelihood to identify phylogenetic signal. As a non-parametric complement, we computed Spearman’s rank correlation between genome size and repeat genomic coverage (Mb). Both analyses were repeated excluding *Belonocnema kinseyi* as its substantially larger genome size relative to the other assemblies could disproportionately influence results.

## Results and Discussion

### Genome assembly metrics

PacBio HiFi sequencing yielded a highly contiguous genome for *Q. erythrinae*. From one Revio cell where the barcoded *Q. erythrinae* SMRTBell library represents 1/10th of the sequencing run, 621,619 HiFi reads (9.5 Gb of HiFi data) were obtained, of which 18 (0.00002% of reads) were discarded after filtering with HiFiAdapterFilt. K-mer analysis of the final assembly relative to the PacBio HiFi reads used to create the contig assemblies reported a raw QV score 55.031. The genome coverage was estimated at 22.6× coverage using GenomeScope2. The initial HiFiASM assembly consisted of 41 contigs, L50 of 5, N50 of 32.9 MB, L90 of 5, N90 of 4.2 MB, and a genome size of 399.197 MB (Table 1). The final assembly consists of 41 contigs representing five autosomes, a mitochondrial genome, and one unplaced 16.7 kb contig composed completely of repeat elements as identified by EarlGrey. Scaffolding with HiC further improved the contiguity of the genome, while BlobToolKit identified one scaffold of non-Arthropod origin, mapped to the endosymbiotic bacterium *Wolbachia pipientis*. Upon removal of the endosymbiont, the final genome consisted of six scaffolds totaling 399.201 MB, an N50 of 75.621 MB, L50 of three, N90 of 59.692 MB, and L90 of five (**Figure 2A**). The assembly has a fairly complete Hymenoptera BUSCO, with 91.1% single-copy complete, 2.4% fragmented, 1.23% duplicated, and 8.9% missing using hymenoptera_odb12 (n=5920) (**Figure 2A**). This genome is in parity with other annotated Chalcidoidea genomes in NCBI (e.g. *Nasonia vitripennis* 95.3%, *Copidosoma floridanum* 90.6%, *Phymastichus coffea* 92.7%; *Trichogramma pretiosum* 92.7%). HiC scaffolding revealed five chromosomes (**Figure 2B**). Repetitive elements encompassed 57.2% (229.8 MB) of the *Q. erythrinae* genome. Among these, TEs class I (LTRs, LINEs, and Penelope-Like Elements) represented 19.2% of the genome, while TEs class II (DNA transposons and Rolling Circle Helitrons) represented 11.0%. Additionally, 24.7% of the TEs were unclassified, while other repeats (Simple Repeats, Microsatellites, RNAs) consisted of 2.4%. The *Q. erythrinae* genome has many quality metrics that meet or exceed the standards of the Earth BioGenome Project, such as a contig N50 well over 1 MB (N50 = 40.5 MB), and single-copy complete BUSCOs over 90% (Lawniczak et al., 2022). Additionally, the size of the scaffold genome (399 MB) is well within the range of size of available genomes on NCBI for Eulophidae (172–509MB) and Chalcidoidea (172–1085 MB).

**Table 1.** Assembly statistics of the *Quadrastichus erythrinae* genome contig and scaffold assemblies.

|  | Contig | Scaffold |
| --- | --- | --- |
| Total Fragments | 41 | 6 |
| N50 (MB) | 32.9 | 75.6 |
| L50 | 5 | 3 |
| N90 (MB) | 4.2 | 59.7 |
| L90 | 5 | 3 |
| Total (MB) | 399.2 | 399.2 |

**Figure 2.**
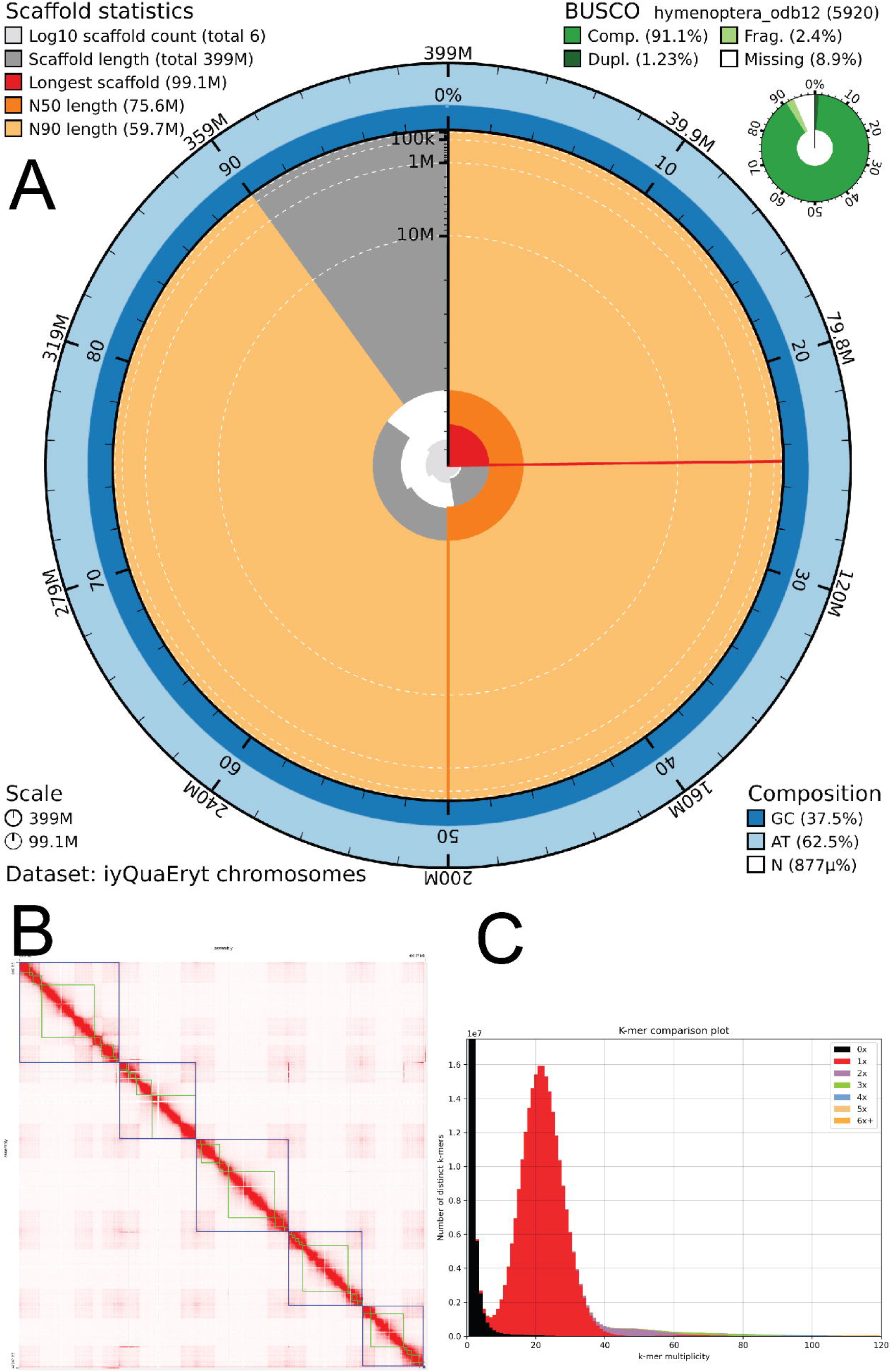
Statistics of the *Quadrastichus erythrinae* assembly. A) Snail plot visualization of the scaffold assembly with BUSCO assessment using Hymenoptera conserved orthologs odb12. B) Hi-C contact map indicates that contigs can be grouped in 5 major scaffolds. C) K-mer spectra plot showing the representation graph of k-mers in the assembly relative to the raw HiFi data.

The mitochondrial genome of *Quadrastichus erythrinae* is 14,607 bp in length and contains the full complement of 13 protein-coding genes, 22 tRNA genes, and both rRNA genes. The arrangement of genes is cox1–cox2–atp8–atp6–cox3–nad3–nad2–rrnS–rrnL–nad1–cytb–nad6–nad4L–nad4–nad5, which differs from the type-5 and type-6 arrangements found in many chalcidoids, including other *Quadrastichus* species (Zhu et al., 2023). In particular, the cox/atp/nad3 block is inverted compared to type-5/6, with type-5/6 showing nad3–cox3–atp6–atp8–cox2–cox1 while the genome of *Q. erythrinae* has cox1–cox2– atp8–atp6–cox3–nad3; the nad5–nad4–nad4L block is also reversed relative to type-5/6, and nad2 is moved from the last position in type-5/6 to an early position in the genome, reflecting extensive mitochondrial gene rearrangement.

The *Wolbachia* genome wQua is 1,346,620bp, with a BUSCO score of 98.0% single-copy complete, 0.3% fragmented, 1.7% duplicated, and 0% missing using rickettsiales_odb12 (n=345). It belongs to Supergroup A, which is commonly found among terrestrial insects including Hymenoptera (Vancaester and Blaxter, 2023).

### Chromosome-level syntenic conservation

Whole-genome alignments indicate high syntenic conservation between *Q. erythrinae* and other hymenopteran genomes despite considerable variation in haploid chromosome counts and assembly contiguity across taxa (**Figure 3**). Reciprocal best-hit comparisons across all annotated genes reveal the highest degree of orthology between the two chalcidoid species *Q. erythrinae* and *N. vitripennis* (*n* = 7,144), intermediate counts in other comparisons, and the lowest between *B. kinseyi* and *D. longicaudata* (*n* = 691) after filtering for syntenic blocks containing ≥10 gene anchors (**Figure 3A**). Gene order between *Q. erythrinae* and *N. vitripennis* is largely collinear (**Figure 3B**), consistent with extensive conservation of chromosomal architecture within Chalcidoidea.

**Figure 3.**
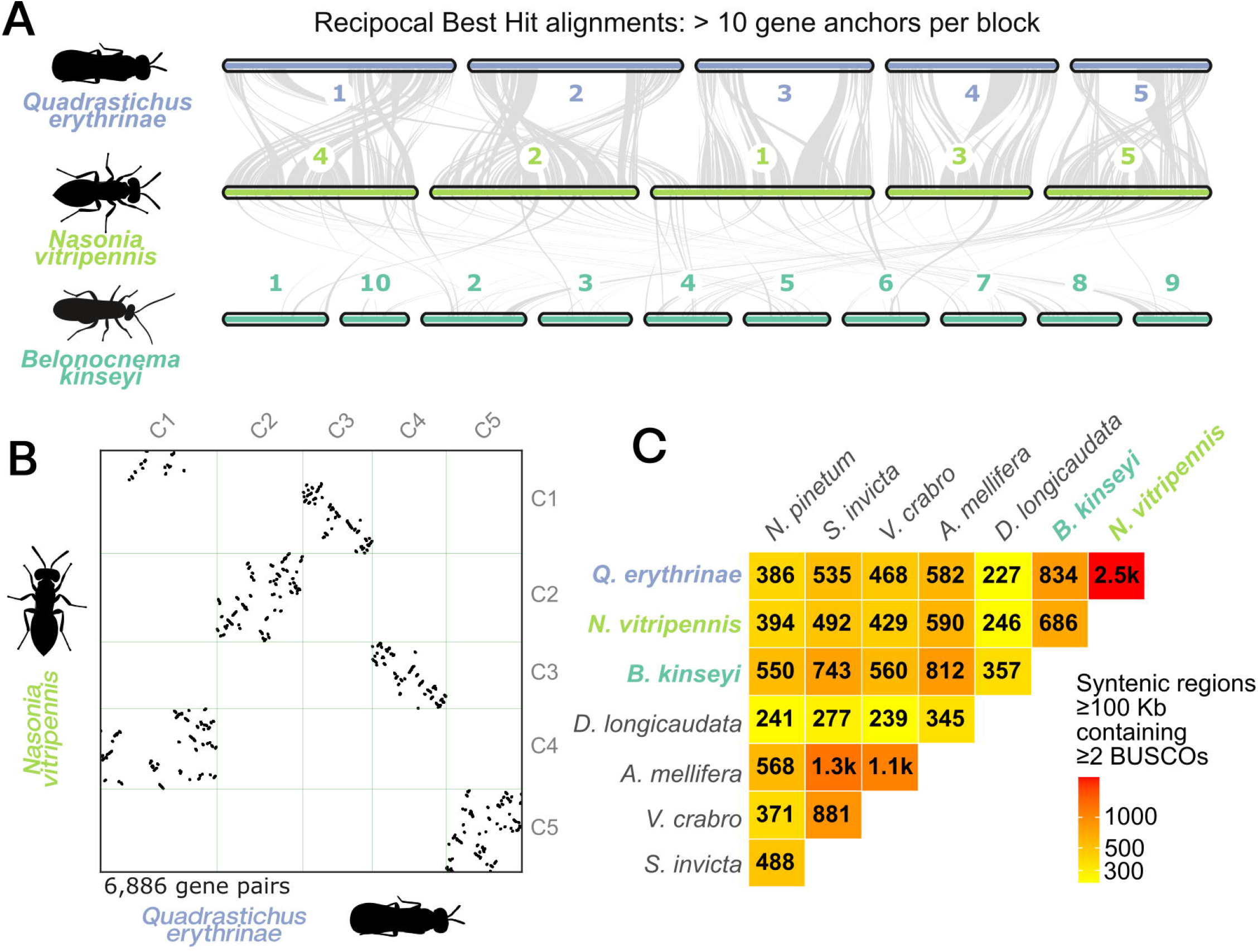
Gene-based synteny across eight hymenopteran genomes. **(A)** JCVI karyoplot showing reciprocal best-hit gene alignments across chromosomes inferred from the full genome annotation. Chromosome lengths reflect gene order only and do not represent physical sizes. **(B)** Pairwise dot plot showing collinearity between *Quadrastichus erythrinae* and *Nasonia vitripennis* based on reciprocal best-hit alignments. (**C**) Pairwise counts of conserved macro-syntenic blocks inferred with chromsyn using BUSCO-defined anchors, filtered for collinear blocks ≥100 Kb containing ≥2 genes.

At the level of deeply conserved genes, BUSCO counts on chromosome-scale scaffolds differed modestly between Proctotrupomorpha (*Q*.*erythrinae, N*.*vintripennis, B*.*kinseyi*) (mean = 5,365, range = 5,234– 5,543) and other lineages (mean = 5,795, range = 5,752–5,826), with overlapping ranges and only a marginally significant difference in mean values (two-sample t-test p-value = 0.04). This pattern indicates broadly comparable recovery of deeply conserved single-copy orthologs across assemblies, consistent with BUSCO scores as a measure of expected gene content completeness rather than a sensitive indicator of lineage-specific gene gain or loss (Waterhouse et al. 2018). Even across deep evolutionary distances in Hymenoptera (∼250 My divergence; (Blaimer et al. 2023)) we recover numerous macro-syntenic blocks ≥100 Kb containing at least two BUSCO genes, reflecting persistence of large conserved genomic segments despite lineage-specific rearrangements (**Figure 3C**). The greatest number of conserved BUSCO-anchored syntenic blocks is between the two chalcidoids *Q. erythrinae* and *N. vitripennis* (n = 2,482), whereas the fewest occur between *Q. erythrinae* and the braconid *Diachasmimorpha longicaudata* (n = 227). This pattern reflects known heterogeneity in chromosome number evolution across Hymenoptera, with broad karyotypic diversity involving frequent rearrangements accompanying speciation and life-history shifts (Gokhman 2022).

### Repeat evolution

Repeat composition varied substantially across the sampled hymenopteran genomes (**Figure 4**). Notably, no short interspersed nuclear elements (SINEs) were detected on chromosome-scale scaffolds in any assembly, consistent with previous reports that SINEs comprise a negligible fraction (<1 %) of transposable element content in Hymenoptera relative to other insect orders (Petersen et al. 2019). Despite relatively similar gene counts across species (ranging from 33,367 in *Apis mellifera* to 44,526 in *Solenopsis invicta*), genome size varied substantially, leading us to test whether differences in repeat abundance explain this interspecific variation. Across the seven species excluding the exceptionally large-genome outlier *B. kinseyi* (1.54 Gb, Table 2), phylogenetically corrected regression revealed that total repeat content is a strong predictor of genome size (β = 0.32 ± 0.086, p = 0.014, λ = 0.0), explaining 73.6 % of variance in genome size (**Figure 4D**). A non-parametric Spearman correlation corroborated this association (ρ = 0.89, p = 0.012), further supporting a positive relationship between repeats and genome size (**Figure 4D**). These associations remained significant with inclusion of *B. kinseyi* (PGLS p < 0.01; Spearman’s ρ = 0.93; **Figure 4C**). These patterns align with patterns observed broadly across insects, where repetitive sequence content often underlies genome size variation (Cong et al. 2022; Cook et al. 2025) and contributes to the C-value enigma by driving genome size expansions independent of coding content (Gregory 2001). Interestingly, despite *Q. erythrinae* having a much smaller genome than *B. kinseyi*, both distantly related gall inducers harbor large proportions of unclassified TEs, accounting for 24.7% and 52.8% of their genomes, respectively (**Figure 4A**). This contrasts with the more closely related chalcidoid parasitoid *N. vitripennis*, which also has a small genome but a much lower proportion of unclassified TEs (13.5%). Although it is premature to infer convergent genomic patterns among phytophagous lineages within Proctotrupomorpha, future characterization of these unclassified TEs across broader taxonomic sampling within the group will be important for determining whether genome expansion is associated with the transition to phytophagy. We note that *S. invicta* also exhibits an increase in distinct TE families (Fig. 4A), highlighting that lineage-specific TE proliferation occurs across Hymenoptera and emphasizing the need for broader sampling to disentangle ecological from lineage-specific effects.

**Figure 4.**
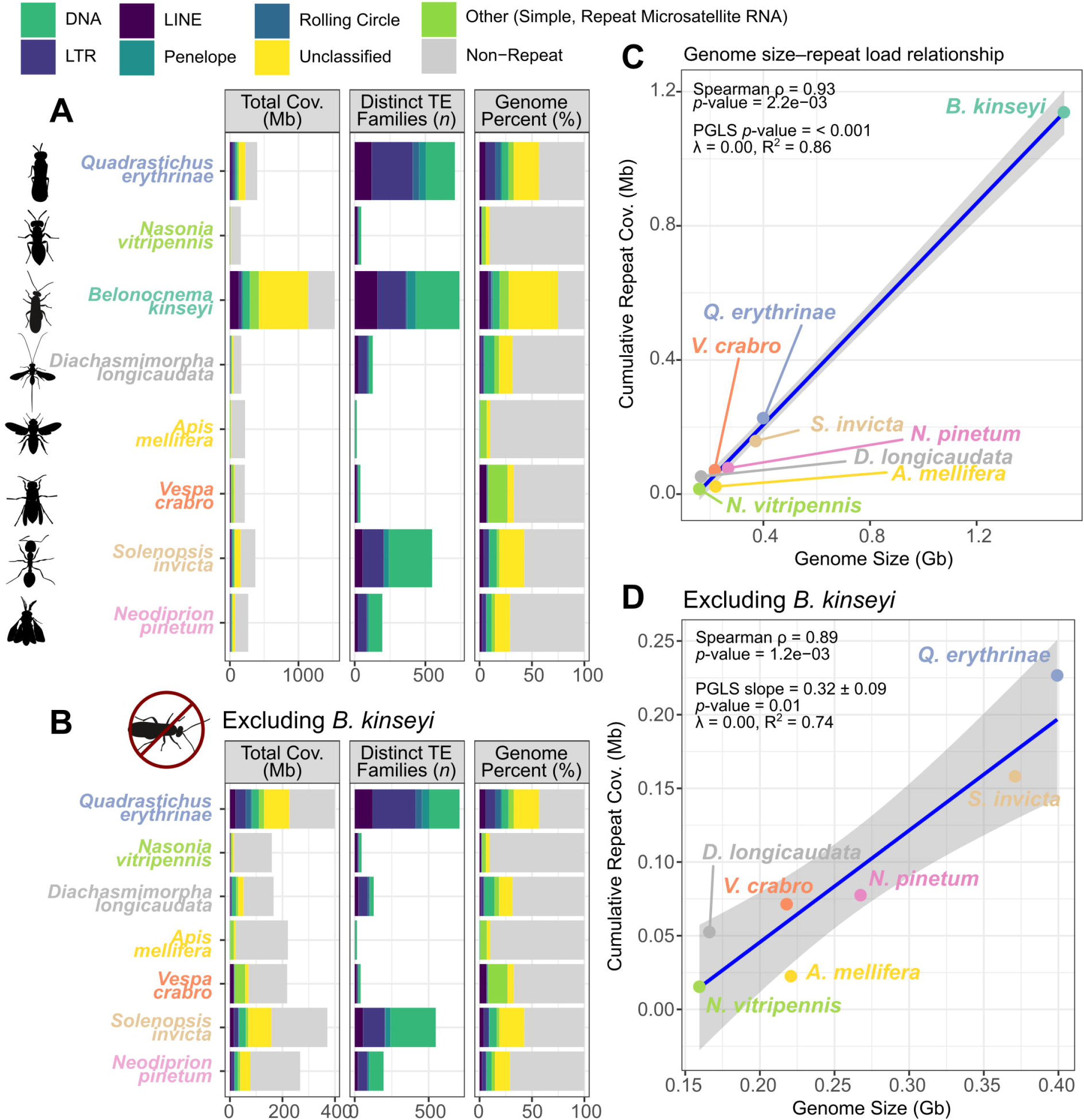
Repeat landscape across hymenopteran genomes. (**A**) Total repetitive DNA content was quantified for eight chromosome-level hymenopteran assemblies using EarlGrey and summarized by repeat subclass. Facets show total cumulative genomic coverage (Mb), number of distinct transposable element (TE) families, and percent of the genome occupied by each repeat subclass. “Non-Repeat”, “Unclassified”, and “Other” categories were excluded from TE family count visualizations (**B**) Same as above, except excluding the large-genome outlier *Belonocnema kinseyi* (bottom). (**C**) Association between genome size (Gb) and cumulative repeat coverage (Gb) with PGLS regression and Spearman correlation results annotated. (**D**) Same analysis excluding *B. kinseyi*.

**Table 2.**
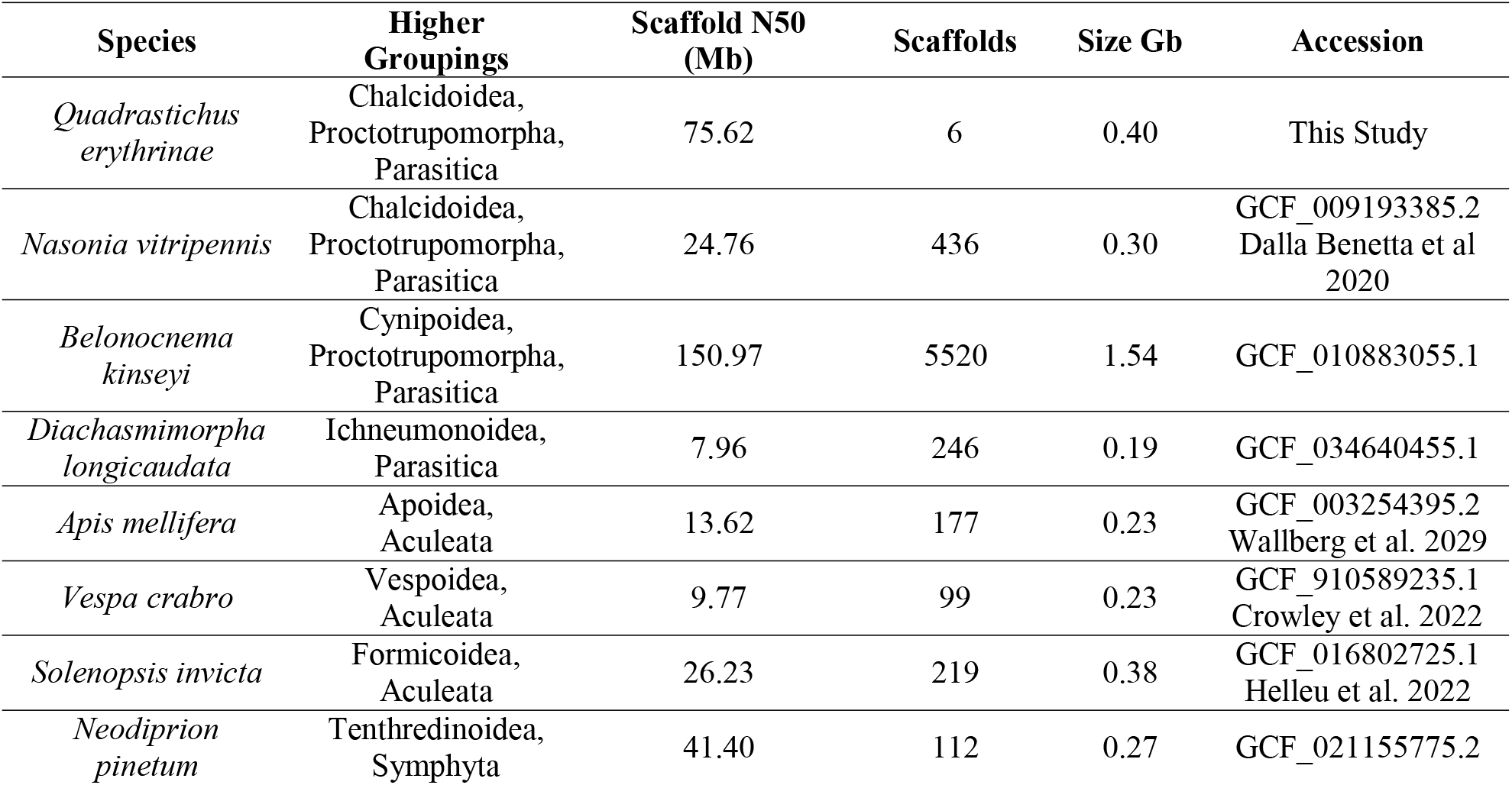
Assembly statistics and accession information for the eight hymenopteran genomes analyzed in this study.

| Species | Higher Groupings | Scaffold N50 (Mb) | Scaffolds | Size Gb | Accession |
| --- | --- | --- | --- | --- | --- |
| *Quadrastichus erythrinae* | Chalcidoidea, Proctotrupomorpha, Parasitica | 75.62 | 6 | 0.40 | This Study |
| *Nasonia vitripennis* | Chalcidoidea, Proctotrupomorpha, Parasitica | 24.76 | 436 | 0.30 | GCF_009193385.2 Dalla Benetta et al 2020 |
| *Belonocnema kinseyi* | Cynipoidea, Proctotrupomorpha, Parasitica | 150.97 | 5520 | 1.54 | GCF_010883055.1 |
| *Diachasmimorpha longicaudata* | Ichneumonoidea, Parasitica | 7.96 | 246 | 0.19 | GCF_034640455.1 |
| *Apis mellifera* | Apoidea, Aculeata | 13.62 | 177 | 0.23 | GCF_003254395.2 Wallberg et al. 2029 |
| *Vespa crabro* | Vespoidea, Aculeata | 9.77 | 99 | 0.23 | GCF_910589235.1 Crowley et al. 2022 |
| *Solenopsis invicta* | Formicoidea, Aculeata | 26.23 | 219 | 0.38 | GCF_016802725.1 Helleu et al. 2022 |
| *Neodiprion pinetum* | Tenthredinoidea, Symphyta | 41.40 | 112 | 0.27 | GCF_021155775.2 |

### Conclusion

The *Q. erythrinae* reference genome, generated from a single female measuring <2 mm with <100ng of input DNA for library preparation and occupying approximately one-tenth of a PacBio HiFi SMRT Cell, represents a major advance as the first genome for a phytophagous eulophid and the first chromosome-scale assembly for the genus *Quadrastichus*. Importantly, this demonstrates that high-quality, chromosome-level assemblies can be obtained from extremely small insects without whole-genome amplification, defining a practical lower bound for direct long-read sequencing of minute taxa. Given current PacBio Revio pricing, the HiFi sequencing for this genome corresponds to only ∼$100 USD in marginal sequencing cost, underscoring the increasing feasibility of low-cost *de novo* genome assembly for tiny hymenopterans, even while acknowledging additional costs associated with sample preparation and library construction. As only the fifth genome available for the hyperdiverse subfamily Tetrastichinae—and only the second at chromosome scale—this resource expands genomic representation for Chalcidoidea. Comparative analyses reveal strong macro-syntenic conservation across Hymenoptera despite extensive karyotypic variation, highlighting the utility of this assembly for evolutionary and phylogenomic studies. Patterns of repeat content further indicate that genome size evolution in Hymenoptera is largely driven by repetitive elements, with *Q. erythrinae* exhibiting relatively low repeat divergence compared to other lineages. This high-quality assembly facilitates population genomic, diagnostic, and phylogenetic analyses, including the identification of loci underlying cryptic species boundaries and traits such as gall induction. In addition, recovery of a complete *Wolbachia* genome directly from host sequencing data underscores the value of chromosome-level assemblies for integrative host–symbiont research. Together, these resources establish the *Q. erythrinae* genome as a foundational genomic model for both applied pest management and comparative hymenopteran genomics.

## Data Availability

Custom scripts, plotting input files, and alignments used for chromosome-scale synteny and repeat analyses are available at https://github.com/merondun/hymenopteran_alignments and are permanently archived on Zenodo (DOI: https://doi.org/10.5281/zenodo.18157483). The primary and alternate genome assemblies are hosted at the National Center for Biotechnology Information (NCBI) under BioProject Accessions PRJNA1404342 and PRJNA1404344 respectively. The sample used for the contig assembly is described under BioSample SAMN54712453 and registered under the Darwin Tree of Life ID iyQuaEryt1. Raw read data was submitted under SRA accessions: SRR36875831-SRR36875835.

## Acknowledgments

YMZ is supported by Oak Ridge Institute for Science and Education (ORISE) fellowship project number 60-2040-2-011. This research used resources provided by the USDA-ARS project number 2040-30400-003-000D, USDA-ARS SCINet project numbers 0201-88888-003-000D and 0201-88888-002-000D. This research was performed as part of the USDA-ARS Ag100Pest Initiative.

## References

Astashyn A, Tvedte ES, Sweeney D, Sapojnikov V, Bouk N, Joukov V, Mozes E, Strope PK, Sylla PM, Wagner L. 2024. Rapid and sensitive detection of genome contamination at scale with FCS-GX. Genome Biol. 25:60. doi:10.1186/s13059-024-03198-7.

Baril T, Galbraith J, Hayward A. 2024. Earl Grey: a fully automated user-friendly transposable element annotation and analysis pipeline. Mol Biol Evol. 41(4):msae068. doi:10.1093/molbev/msae068.

Bates D, Mächler M, Bolker B, Walker S. 2015. Fitting Linear Mixed-Effects Models Using lme4. J Stat Softw. 67(1). doi:10.18637/jss.v067.i01.

Bernt M, Donath A, Jühling F, Externbrink F, Florentz C, Fritzsch G, Pütz J, Middendorf M, Stadler PF. 2013. MITOS: improved de novo metazoan mitochondrial genome annotation. Mol Phylogenet Evol. 69(2):313–319. doi:10.1016/j.ympev.2012.08.023.

Blaimer BB, Brady SG, Buffington ML, Gates MW, Kula RR, et al. 2023. Key innovations and the diversification of Hymenoptera. Nat Commun. 14(1):1212. doi:10.1038/s41467-023-36868-4.

Buchfink B, Reuter K, Drost HG. 2021. Sensitive protein alignments at tree-of-life scale using DIAMOND. Nat Methods. 18(4):366–368. doi:10.1038/s41592-021-01101-x.

Camacho C, Coulouris G, Avagyan V, Ma N, Papadopoulos J, Bealer K, Madden TL. 2009. BLAST+: architecture and applications. BMC Bioinformatics. 10:421. doi:10.1186/1471-2105-10-421.

Challis R, Richards E, Rajan J, Cochrane G, Blaxter M. 2020. BlobToolKit: interactive quality assessment of genome assemblies. G3 (Bethesda). 10(4):1361–1374. doi:10.1534/g3.119.400908.

Cheng H, Concepcion GT, Feng X, Zhang H, Li H. 2021. Haplotype-resolved de novo assembly using phased assembly graphs with hifiasm. Nat Methods. 18(2):170–175. doi:10.1038/s41592-020-01056-5.

Cheng H, Jarvis ED, Fedrigo O, Koepfli KP, Urban L, Gemmell NJ, Li H. 2022. Haplotype-resolved assembly of diploid genomes without parental data. Nat Biotechnol. 40(9):1332–1335. doi: 10.1038/s41587-022-01261-x.

Cong Y, Wang J, Lin Y, Chen H, Li Y, Zhang M, Zhang H, Li Z, Li X, Yu W, et al. 2022. Transposons and non-coding regions drive the intrafamily differences of genome size in insects. iScience. 25(9):104873. doi:10.1016/j.isci.2022.104873.

Cook HL, Sproul JS, Murray EA, Bossert S. 2025. A comparative analysis of transposable element diversity and evolution across 75 bee genomes. BMC Genomics. 26(1):1000. doi:10.1186/s12864-025-12190-9.

Crowley LM, University of Oxford and Wytham Woods Genome Acquisition Lab, Darwin Tree of Life Barcoding collective, Wellcome Sanger Institute Tree of Life programme, Wellcome Sanger Institute Scientific Operations: DNA Pipelines collective, et al. The genome sequence of the European hornet, Vespa crabro Linnaeus, 1758. Wellcome Open Res 2022, 7:27 doi:10.12688/wellcomeopenres.17546.1

Dalla Benetta E, Antoshechkin I, Yang T, Nguyen HQ, Ferree PM, Akbari OS. 2020 Genome elimination mediated by gene expression from a selfish chromosome. Sci. adv. 6(14):eaaz9808. Doi: 10.1126/sciadv.aaz9808

DeAngelis MM, Wang DG, Hawkins TL. 1995. Solid-phase reversible immobilization for the isolation of PCR products. Nucleic Acids Res. 23(22):4742–4743. doi:10.1093/nar/23.22.4742.

Edwards RJ, Dong C, Park RF, Tobias PA. 2022. A phased chromosome-level genome and full mitochondrial sequence for the dikaryotic myrtle rust pathogen, Austropuccinia psidii. bioRxiv. doi:10.1101/2022.04.22.489119.

Frith MC, Hamada M, Horton P. 2010. Parameters for accurate genome alignment. BMC Bioinformatics. 11:80. doi:10.1186/1471-2105-11-80.

Gokhman VE. 2022. Comparative Karyotype Analysis of Parasitoid Hymenoptera (Insecta): Major Approaches, Techniques, and Results. Genes (Basel). 13(5):751. doi:10.3390/genes13050751.

Gregory TR. 2001. Coincidence, coevolution, or causation? DNA content, cell size, and the C-value enigma. Biol Rev Camb Philos Soc. 76(1):65–101. doi:10.1017/S1464793100005595.

Guan D, McCarthy SA, Wood J, Howe K, Wang Y, Durbin R. 2020. Identifying and removing haplotypic duplication in primary genome assemblies. Bioinformatics. 36(9):2896–2898. doi:10.1093/bioinformatics/btaa025.

Haas BJ, Papanicolaou A, Yassour M, Grabherr M, Blood PD, Bowden J, Couger MB, Eccles D, Li B, Lieber M, et al. 2013. De novo transcript sequence reconstruction from RNA-seq using the Trinity platform for reference generation and analysis. Nat Protoc. 8(8):1494–1512. doi:10.1038/nprot.2013.084.

Helleu Q, Roux C, Ross KG, Keller L. 2022. Radiation and hybridization underpin the spread of the fire ant social supergene. Proc. Natl. Acad. Sci. USA. 119(34):e2201040119. doi: 10.1073/pnas.2201040119

[Jacques Dainat] NBISweden/AGAT: AGAT v1.5.1 [software]. 2025. Zenodo. doi:10.5281/zenodo.3552717.

Kärrnäs E, Hansson C, Wahlberg N. 2025. Large-scale DNA barcoding reveals cryptic diversity in eulophid wasps (Hymenoptera, Chalcidoidea, Eulophidae). Arthropod Syst Phylogeny. 83:415–425. doi:10.3897/asp.83.e153226.

Kaufman LV, Yalemar J, Wright MG. 2020. Classical biological control of the erythrina gall wasp, Quadrastichus erythrinae, in Hawaii: conserving an endangered habitat. Biol Control. 142:104161. doi: 10.1016/j.biocontrol.2019.104161.

Kim IK, Delvare G, La Salle J. 2004. A new species of Quadrastichus (Hymenoptera: Eulophidae): a gall-inducing pest on Erythrina (Fabaceae). J Hymenopt Res. 13(2):243–249.

Lawniczak MKN, Durbin R, Flicek P, Lindblad-Toh K, Wei X, Archibald JM, Baker WJ, Belov K, Blaxter ML, Marques Bonet T, et al. 2022. Standards recommendations for the Earth BioGenome Project. Proc Natl Acad Sci U S A. 119(4):e2115639118. doi:10.1073/pnas.2115639118.

Lawrence M, Huber W, Pagès H, Aboyoun P, Carlson M, Gentleman R, Morgan MT, Carey VJ. 2013. Software for computing and annotating genomic ranges. PLoS Comput Biol. 9(8):e1003118. doi:10.1371/journal.pcbi.1003118.

Lenth R, Piaskowski J. emmeans: Estimated Marginal Means, aka Least-Squares Means. R package. 2025.

Li H. 2018. Minimap2: pairwise alignment for nucleotide sequences. Bioinformatics. 34(18):3094–3100. doi:10.1093/bioinformatics/bty191.

Lin SF, Tung GS, Yang MM. 2021. The Erythrina gall wasp Quadrastichus erythrinae (Insecta: Hymenoptera: Eulophidae): invasion history, ecology, infestation and management. Forests. 12(7):948. doi:10.3390/f12070948.

Manni M, Berkeley MR, Seppey M, Simão FA, Zdobnov EM. 2021. BUSCO Update: Novel and Streamlined Workflows along with Broader and Deeper Phylogenetic Coverage for Scoring of Eukaryotic, Prokaryotic, and Viral Genomes. Mol Biol Evol. 38(10):4647–4654. doi:10.1093/molbev/msab199.

Orme D, Freckleton R, Thomas G, Petzoldt T, Fritz S. caper: Comparative Analyses of Phylogenetics and Evolution in R. R package. 2025.

Petersen M, Armisen D, Gibbs RA, Hering L, Khila A, Mayer G, Richards S, Niehuis O, Misof B. 2019. Diversity and evolution of the transposable element repertoire in arthropods with particular reference to insects. BMC Evol Biol. 19(1):11. doi:10.1186/s12862-018-1324-9.

R Core Team. R: A language and environment for statistical computing [software]. 2017. R Foundation for Statistical Computing, Vienna, Austria.

Rasplus JY, Basso C, Delvare G, Fusu L, Gumovsky A, Huber JT, Polaszek A, Schmidt S, Murphy N, Butterill P, et al. 2020. A first phylogenomic hypothesis for Eulophidae (Hymenoptera, Chalcidoidea). J Nat Hist. 54(9-10):597–609. doi: 10.1080/00222933.2020.1762941.

Seemann T. 2014. Prokka: rapid prokaryotic genome annotation. Bioinformatics. 30(14):2068–2069. doi:10.1093/bioinformatics/btu153.

Sim SB, Corpuz RL, Simmonds TJ, Geib SM. 2022. HiFiAdapterFilt: a memory-efficient read processing pipeline for PacBio HiFi reads. BMC Genomics. 23(1):157. doi:10.1186/s12864-022-08375-1.

Sproul JS, Hotaling S, Heckenhauer J, Powell A, Larracuente AM, Pauls SU, Kelley JL. 2023. Analyses of 600+ insect genomes reveal repetitive element dynamics and highlight biodiversity-scale repeat annotation challenges. Genome Res. 33(10):1708–1717. doi:10.1101/gr.277387.122.

Tang H, DeBarry JD, Tan X, Wang X, Moore D, He Z, Luo M, Lee TH, Jin H, Kudrna D, et al. 2024. JCVI: A versatile toolkit for comparative genomics analysis. iMeta. 3(4):e211. doi:10.1002/imt2.211.

Tegenfeldt F, Kuznetsov D, Manni M, Berkeley M, Zdobnov EM, Kriventseva EV. 2025. OrthoDB and BUSCO update: annotation of orthologs with wider sampling of genomes. Nucleic Acids Res. 53(D1):D516–D522. doi:10.1093/nar/gkae987.

[UCD Community]. Universal Chalcidoidea Database. 2023. [accessed 2025 Nov 21]. https://ucd.chalcid.org.

Uliano-Silva M, Ferreira JGRN, Krasheninnikova K, Darwin Tree of Life Consortium, Blaxter M, et al. 2023. MitoHiFi: a python pipeline for mitochondrial genome assembly from PacBio high-fidelity reads. BMC Bioinformatics. 24(1):288. doi:10.1186/s12859-023-05385-y.

Vancaester E, Blaxter M. 2023. Phylogenomic analysis of Wolbachia genomes from the Darwin Tree of Life biodiversity genomics project. PLoS Biol. 21(8):e3001972. doi:10.1371/journal.pbio.3001972.

Vasimuddin M, Misra S, Li H, Aluru S. 2019. Efficient architecture-aware acceleration of BWA-MEM for multicore systems. In: 2019 IEEE International Parallel and Distributed Processing Symposium (IPDPS). p. 314–324. doi:10.1109/IPDPS.2019.00041.

Wallberg A, Bunikis I, Pettersson OV, Mosbech MB, Childers AK et al. A hybrid de novo genome assembly of the honeybee, Apis mellifera, with chromosome-length scaffolds. BMC Genomics 20, 275 (2019). doi:10.1186/s12864-019-5642-0

Waterhouse RM, Seppey M, Simão FA, Manni M, Ioannidis P, Klioutchnikov G, Kriventseva EV, Zdobnov EM. 2018. BUSCO Applications from Quality Assessments to Gene Prediction and Phylogenomics. Mol Biol Evol. 35(3):543–548. doi:10.1093/molbev/msx319.

Wickham H, Averick M, Bryan J, Chang W, McGowan L, François R, Grolemund G, Hayes A, Henry L, Hester J, et al. 2019. Welcome to the Tidyverse. J Open Source Softw. 4(43):1686. doi:10.21105/joss.01686.

Wilke CO. ggridges: Ridgeline Plots in ‘ggplot2’. R package. 2025.

Zhang YM, Gates MW, Hanson PE, Jansen-González S. 2022. Description of a Neotropical gall inducer on Araceae: Arastichus gen. nov. (Hymenoptera, Eulophidae) and two new species. J Hymenopt Res. 92:145–172. doi: 10.3897/jhr.92.85967.

Zhou C, McCarthy SA, Durbin R. 2023. YaHS: yet another Hi-C scaffolding tool. Bioinformatics. 39(1):btac808. doi:10.1093/bioinformatics/btac808.

Zhu JC, Wang YF, Zhang YM, Wang J, Li F, Liu JX, Xiao JH, Huang DW. 2023. Evolutionary timescale of chalcidoid wasps inferred from over one hundred mitochondrial genomes. Zool Res. 44(3):467–480. doi:10.24272/j.issn.2095-8137.2022.379.

